# The fitness consequences of genetic variation in wild populations of mice

**DOI:** 10.1101/383240

**Authors:** Rowan D.H. Barrett, Stefan Laurent, Ricardo Mallarino, Susanne P. Pfeifer, Charles C.Y. Xu, Matthieu Foll, Kazumasa Wakamatsu, Jonathan S. Duke-Cohan, Jeffrey D. Jensen, Hopi E. Hoekstra

## Abstract

Adaptive evolution can occur when genetic change affects traits subject to natural selection. Although selection is a deterministic process, adaptation can be difficult to predict in finite populations because the functional connections between genotype, phenotype, and fitness are complex. Here, we make these connections using a combination of field and laboratory experiments. We conduct a large-scale manipulative field experiment with wild populations of deer mice in distinct habitats to directly estimate natural selection on pigmentation traits and next test whether this selection drives changes in allele frequency at an underlying pigment locus. We find that divergent cryptic phenotypes are repeatedly favoured in each habitat, leaving footprints of selection in the *Agouti* gene. Next, using transgenic experiments in *Mus*, we functionally test one of the *Agouti* mutations associated with survival, a Serine deletion in exon 2, and find that it causes lighter coat colour via changes in its protein binding properties. Finally, we show significant change in the frequency of this mutation in our field experiment. Together, our findings demonstrate how a sequence variant alters phenotype and show the ensuing ecological consequences that drive changes in population allele frequency, thereby revealing the full process of evolution by natural selection.

---

While a growing number of genomic studies have pinpointed genes that contribute to phenotypic evolution^1-7^, often the ecological mechanisms driving trait evolution remain untested. On the other hand, many elegant field studies have documented the action of natural selection on traits^8-14^, but the underlying molecular mechanisms are unclear. Here, we combine a large-scale manipulative field experiment with laboratory-based genetic and biochemical tests to identify both the ecological and molecular mechanisms that cause trait adaptation in a single study of a wild vertebrate. Forging these precise mechanistic connections will aid in predicting the evolutionary consequences of environmental change in natural populations^15,16^.

We took advantage of recently evolved, cryptically coloured populations of deer mice to investigate the evolutionary and genetic consequences of divergent natural selection. The Sand Hills of Nebraska were formed from light-coloured quartz ∼8-10,000 years ago^17^. This dune habitat differs from the surrounding habitat in physical properties, most notably the soil colour^18^ (Fig. 1). Because the Sand Hills are geologically young and ecologically distinct, deer mouse populations inhabiting the area are expected to possess recently evolved and strongly selected adaptations to this novel environment. One striking example of such an adaptation is pigmentation. Classic natural history studies described the dorsal coats of deer mice as strongly correlated with substrate colour, with light mice occupying the newly formed light Sand Hills^19^. The primary hypothesis for this phenotypic change is selection for crypsis against avian predators^19-21^. Pigmentation differences between habitat types are associated with multiple mutations within *Agouti*^22,23^, a locus that mediates the production of yellow pigment (pheomelanin) in vertebrates^24,25^ and deer mice, specifically^21^. Thus, Sand Hills deer mice and the *Agouti* locus are a useful system to directly test both the ecological and functional mechanisms by which specific sequence variants alter phenotype and ultimately fitness.

**Figure 1.**
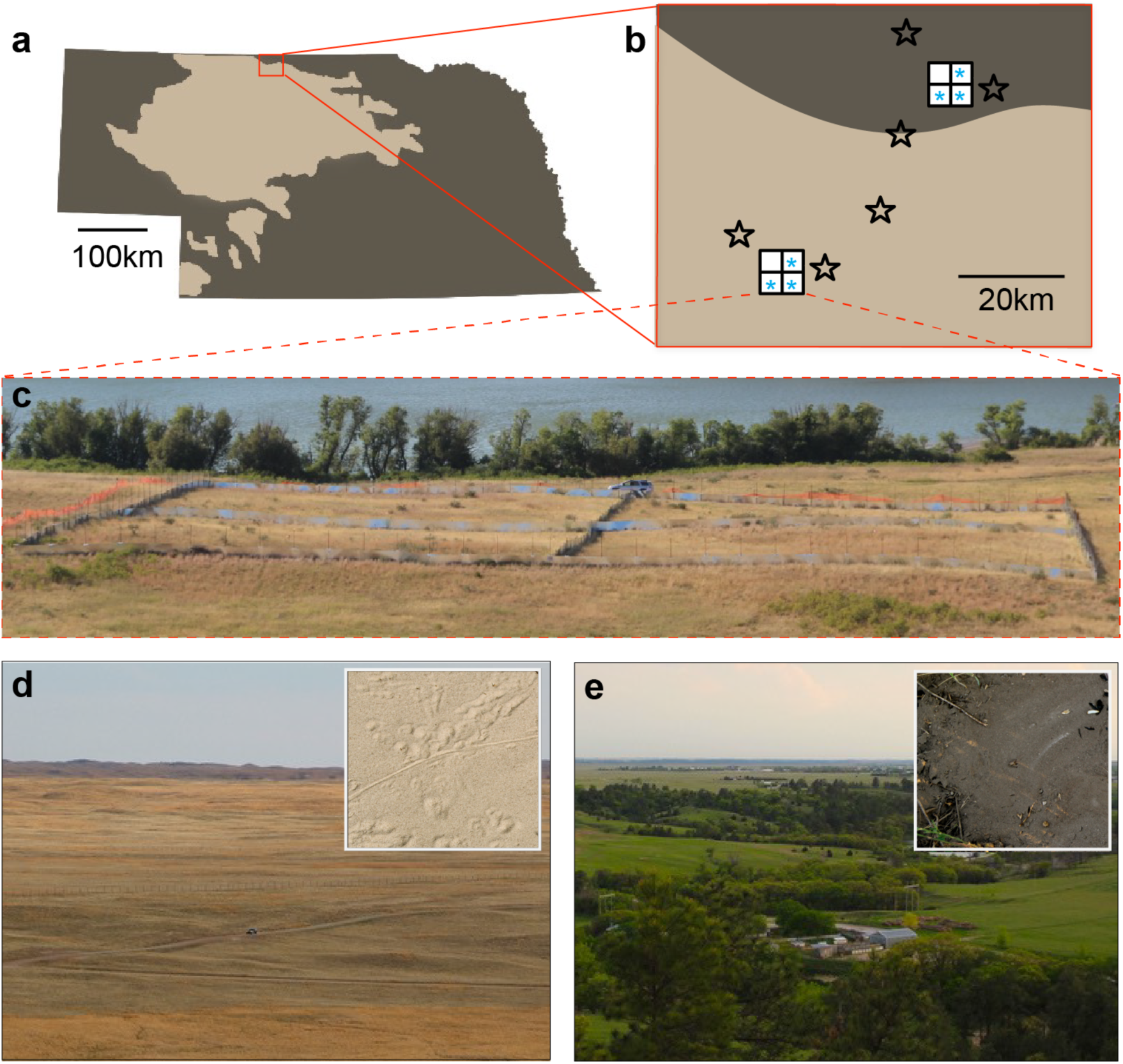
Experimental site and environmental variation in the Sand Hills region of Nebraska. **a**, Map of Nebraska showing the Sand Hills region (light colour) and location of the enclosure experiment. **b**, Map of enclosure sites and sampling locations of wild populations used in the experiment. Location of the replicate enclosures in each habitat type are denoted by white squares. Light blue asterisks indicate the 6 enclosures used. Stars show sampling locations for mice introduced to the enclosures (Extended Data Table 6). **c**, Enclosures are shown at the light site in Sand Hills habitat (truck for scale). **d** and **e**, Typical habitat is shown on the Sand Hills (**d**, light habitat) and off the Sand Hills (**e**, dark habitat); insets show typical soil substrate.

## Divergent selection on pigmentation in experimental enclosures

To explicitly test for divergent natural selection that favours locally adapted pigment phenotypes, we collected 481 wild mice from the ancestral “dark” and derived “light” sites. We then introduced 75-100 individuals in equal proportions based on capture site (i.e., dark versus light) to six 2,500m^2^ field enclosures (three in each habitat) that were devoid of native mice and terrestrial predators, but open to avian predators (Fig. 1; see Methods). Among these founding individuals, we identified significant differences in five pigment traits (dorsal brightness, dorsal chroma, ventral brightness, ventral chroma, and tail pattern) between mice captured at dark versus light sites (all traits: *P* < 0.001, Extended Data Fig. 1). Pigment phenotypes were largely independent, with weak and mostly insignificant correlations among traits (*r*^*2*^<0.06 for all traits, Extended Data Table 1), suggesting that these traits can be subjected to selection independently.

**Table 1.** Standardized linear selection gradients (β) for pigmentation traits.

| Trait | Dark site |  |  | Light site |  |  |
| --- | --- | --- | --- | --- | --- | --- |
|  | Enclosure<br>1 | Enclosure<br>2 | Enclosure<br>3 | Enclosure<br>1 | Enclosure<br>2 | Enclosure<br>3 |
| Dorsal<br>brightness | -0.212<br>(0.289) | -0.411**<br>(0.151) | -0.385**<br>(0.156) | 1.115*<br>(0.830) | 0.649**<br>(0.140) | 0.561<br>(0.463) |
| Dorsal<br>chroma | -0.083<br>(0.156) | 0.250<br>(0.137) | 0.034<br>(0.124) | -0.214<br>(0.362) | 0.069<br>(0.211) | -0.321<br>(0.372) |
| Ventral<br>brightness | -0.151<br>(0.149) | 0.027<br>(0.128) | -0.133<br>(0.129) | 0.246<br>(0.577) | -0.102<br>(0.183) | -0.562<br>(0.772) |
| Ventral<br>chroma | -0.114<br>(0.154) | -0.097<br>(0.144) | 0.300**<br>(0.105) | -0.019<br>(0.487) | -0.229<br>(0.163) | 0.133<br>(0.352) |
| Tail<br>pattern | -0.006<br>(0.183) | -0.133<br>(0.144) | 0.081<br>(0.135) | -0.139<br>(0.721) | 0.343<br>(0.176) | 0.385<br>(0.478) |

Parentheses give 1 standard error. Standard errors and P values determined by 1000 bootstrap replicates. All selection gradients calculated over the first 3 months following introduction to the enclosures. \*: $P < 0.05$ , \*\*: $P < 0.005$ .

Using a mark-recapture approach, we tracked survival of these 481 introduced individuals during five 2-week sampling periods over 14 months, by which time mortality of these mice reached 100% in most enclosures (Fig. 2a,b). Because sampling error is inversely proportional to the number of survivors, we focused our analyses on a comparison between the colonizing populations and survivors present at time point 1 (approximately 3 months after the start of the experiment), when average survival rates were 45%. Regardless of sampling origin or phenotype, the survival rates were twice as high in dark enclosures relative to light enclosures (60% versus 30% at time point 1). Mice introduced to the enclosures that matched the habitat type in which they were originally sampled had significantly greater survival than non-local mice (logistic regression testing the effect of the interaction between mouse source and experimental treatment on survival: *F*_1_ = 19.43, *P* = 1.04e-05; Fig. 2a,b), suggesting local adaptation of populations from each environment type.

**Figure 2.**
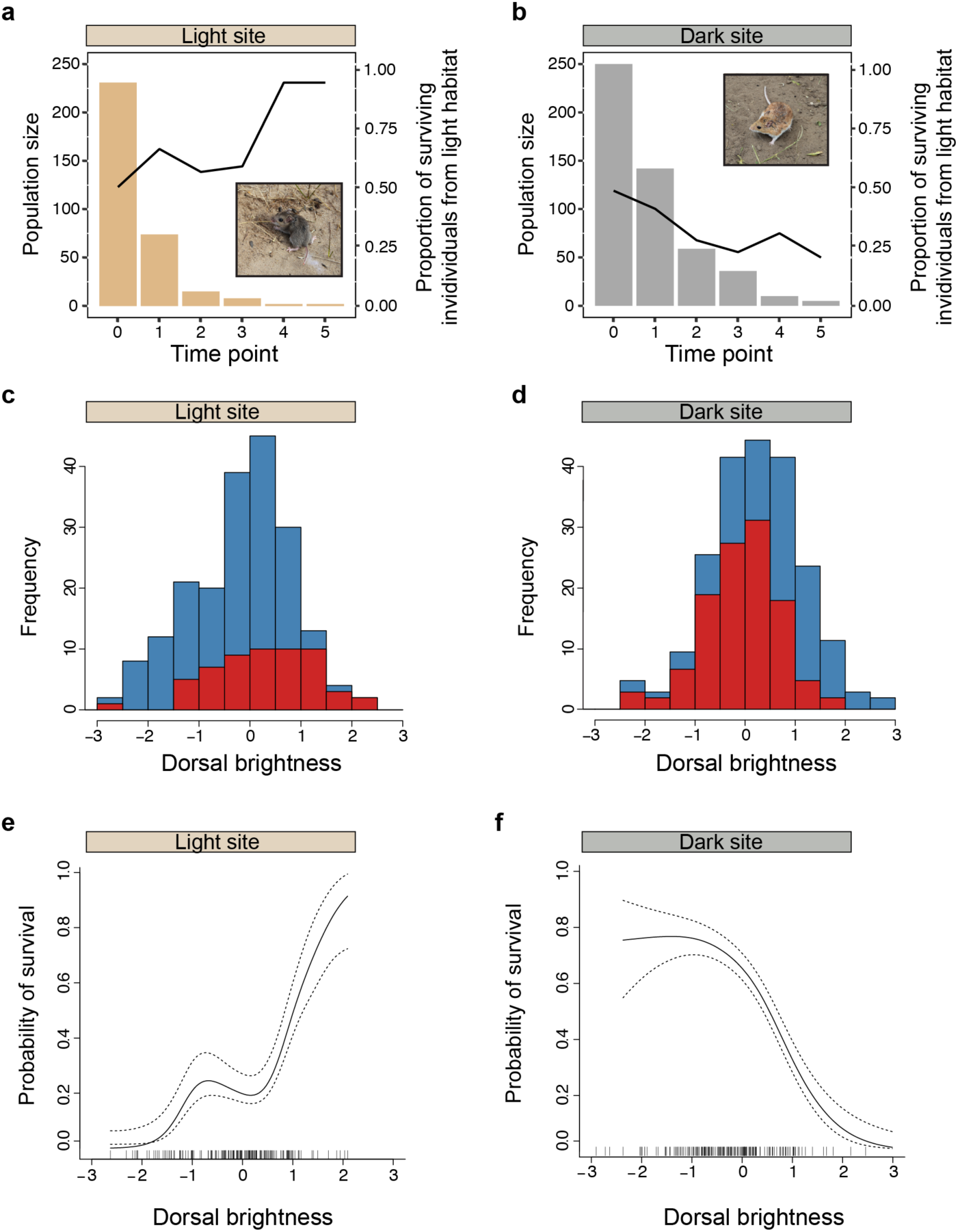
Mortality and phenotypic change in the experimental populations. **a** and **b**, Mortality in pooled enclosures at light (**a**) and dark (**b**) sites over 5 sequential episodes of selection (see Methods). Bars represent the number of surviving individuals (independent of coat colour) at each time point. Black lines represent the proportion of surviving individuals that were originally sampled in light habitat. Conspicuously coloured mice are shown on typical substrate at each experimental site. **c** and **d**, Distributions of dorsal brightness at time point 0 (blue) and time point 1 (red) at the light (**c**) and dark (**d**) sites. **e** and **f**, Visualizations of selection on dorsal brightness at the light (**e**) and dark (**f**) sites. Cubic spline plots are generated from predicted values. The solid lines represent the fitted spline and the dotted lines represent ±1 Bayesian standard error.

To explicitly test if pigmentation may be an important trait contributing to local adaptation, we tested for shifts in the distribution of pigment traits during the experiment. We documented significant selection on pigmentation, primarily manifested through higher survival of mice with locally cryptic dorsal pigmentation (Fig. 2c-f, Table 1). In light enclosures, the surviving mice were, on average, 1.44 times lighter in dorsal colour than the average mouse in the original founding populations, whereas in dark enclosures the average mouse became 1.98 times darker. Linear selection gradients for dorsal brightness were positive in all three light site enclosures and negative in all three dark site enclosures (one-sided t-test of linear selection gradients in light versus dark enclosures: *t* = −6.079, df = 2.518, *P* = 0.015; Table 1). With the exception of ventral chroma in a single enclosure, no significant directional selection was detected on any other trait but dorsal brightness (Table 1). There was also no evidence for quadratic or correlational selection in the data (Extended Data Tables 2,3). Thus, we experimentally demonstrated strong, divergent natural selection on pigmentation between the two environment types, which was confined to a single pigment trait: dorsal brightness.

Our previous work using plasticine model mice to document predation attempts suggests avian predation is high in this region^22^. Moreover, prior experimental work found that owls are highly effective predators of mice and are able to discriminate between different colour morphs even under moonlight conditions^20^. During our field experiment, we observed owls hunting at the experimental sites (8 observations over 70 nights), providing a minimum number of opportunities for owls to predate on mice. Because the enclosures largely exclude other predators, we suggest that the significant association between dorsal brightness and survival is likely driven by higher rates of avian predation on mice with conspicuous pigmentation.

## The genetic consequences of selection on pigmentation

To investigate how the observed selection on dorsal brightness impacts allele frequencies at the *Agouti* locus, we generated polymorphism data with enriched sequencing of a 185-kb region that encompasses *Agouti* and all known regulatory elements as well as in ∼2100 unlinked genome-wide regions averaging 1.5-kb in length (following^26^) to control for demographic effects. In brief, we sequenced all 481 individuals and after filtering, identified 2,442 and 53,507 variable, high-quality sites in or near the *Agouti* gene and genome-wide, respectively. Based on these data, we observed greater changes in allele frequency at *Agouti* over time in the light than dark enclosures, consistent with higher mortality in light enclosures (One-sided ks-test: *D* = 0.143, *P* = 2.2×10^−6^; Extended Data Fig. 2a).

To determine whether the changes in allele frequency at *Agouti* are best explained by selection or neutrality (i.e., genetic drift), we calculated, for every *Agouti* variant site independently, the probability that the distribution of genotype frequencies observed in the survivors is a random sample from the distribution observed in the initial population (see Methods). A large number of SNPs (light enclosures = 353, dark enclosures = 549) showed genotype frequencies in the survivors that cannot be explained by random sampling alone (Fig. 3a, b). To account for the large number of tests involved, we next used a resampling procedure to determine how many SNPs would be expected to show significant change by chance alone. In the light enclosures, the patterns of allele frequency change at *Agouti* SNPs could not be distinguished from neutrality (Fig. 3c), likely due to reduced statistical power caused by low numbers of surviving individuals. By contrast, in the dark enclosures, our results reject the null hypothesis, suggesting that the number of significant changes in allele frequency is incompatible with a neutral model (Fig. 3d). Therefore, in the dark enclosures, we find allele frequency change at the *Agouti* locus that is consistent with the action of selection, and thus patterns at the genetic level parallel the change observed at the phenotypic level.

**Figure 3.**
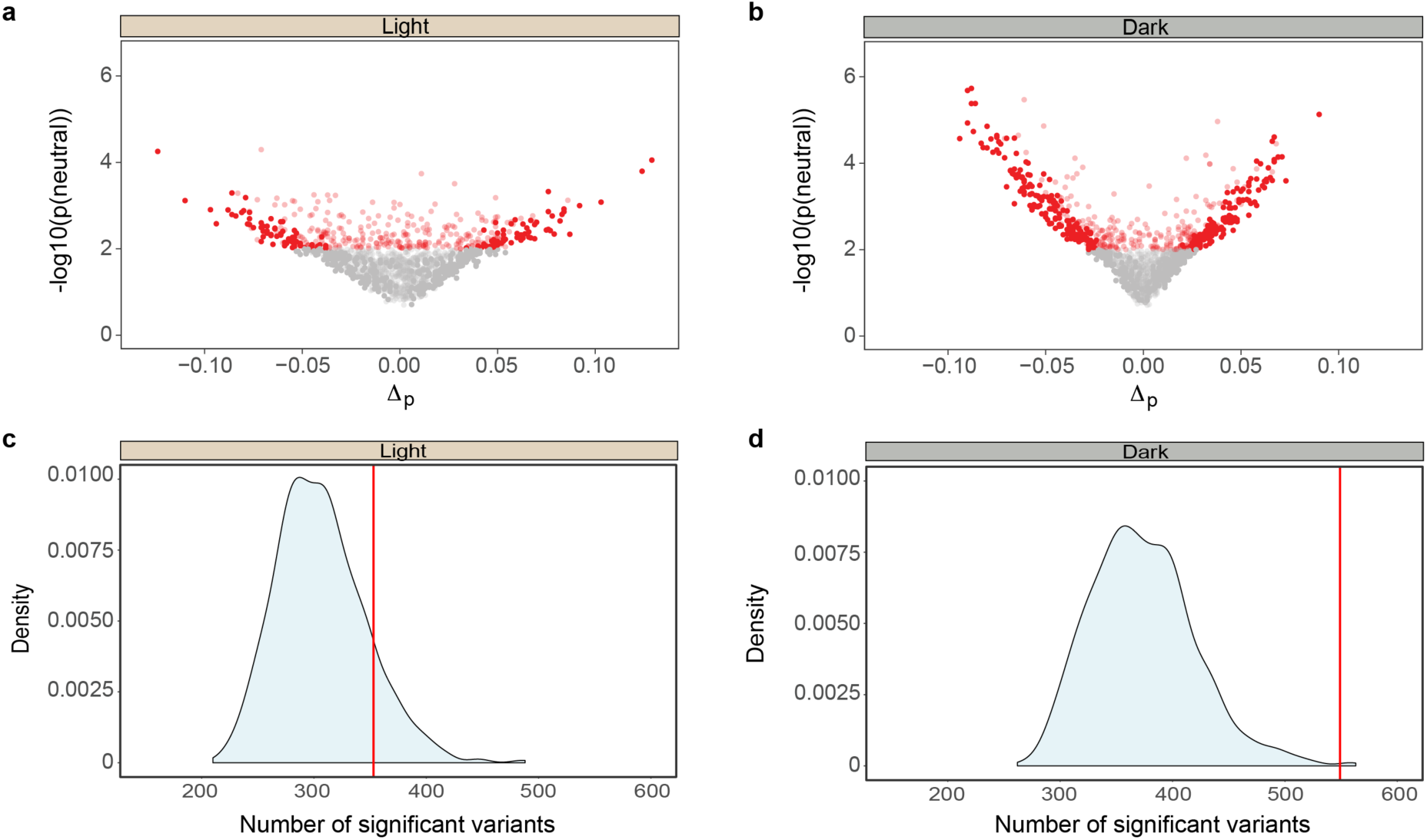
Allele frequency change at the *Agouti* locus. **a** and **b,** Allele frequency change from mortality during the experiment in the pooled light (**a**) and dark (**b**) enclosures. The x-axis represents the change in allele frequency between initial colonizing populations and survivors sampled after 3 months. The y-axis represents the probability of the distribution of genotype frequencies observed in the survivors assuming a neutral model. All red points are significant at the 1% level: light red points are significant due to a bias in the observed proportion of heterozygotes, whereas dark red points exhibit a bias in the observed number of homozygotes. **c** and **d**, Null distributions of the number of sites expected to show significant allele frequency change at the 1% level in the pooled light (**c**) and dark (**d**) enclosures. Vertical red lines represent the observed number of sites with significant allele frequency change.

Because within a single generation there is no recombination between loci and therefore SNPs are not independent, we further tested whether the large number of sites with significant allele frequency change in the dark enclosures could be explained by correlated responses at loci linked to a SNP under selection (see Methods). Based on our phenotypic selection results, we have the *a priori* hypothesis that SNPs associated with dorsal brightness should be experiencing direct selection. Thus, for each of 31 *Agouti* SNPs associated with dorsal brightness^23^, we compared genotype frequencies under a model with and without selection (see Methods). Of these, we show 7 SNPs, including 6 SNPs in or near regulatory regions of *Agouti* and a single amino acid deletion of Serine at amino acid position 48 in exon 2 (ΔSer), whose allele frequency change could not be explained by random sampling alone (Fig. 4a; Extended Data Table 4). Four of these 7 SNPs also had high levels of differentiation between mice captured from light and dark habitats (Fig. 4b; Extended Data Table 4). In addition, one regulatory SNP and the ΔSer previously had been associated with signals of historical positive selection in the Sand Hills populations^22,23^.

**Figure 4.**
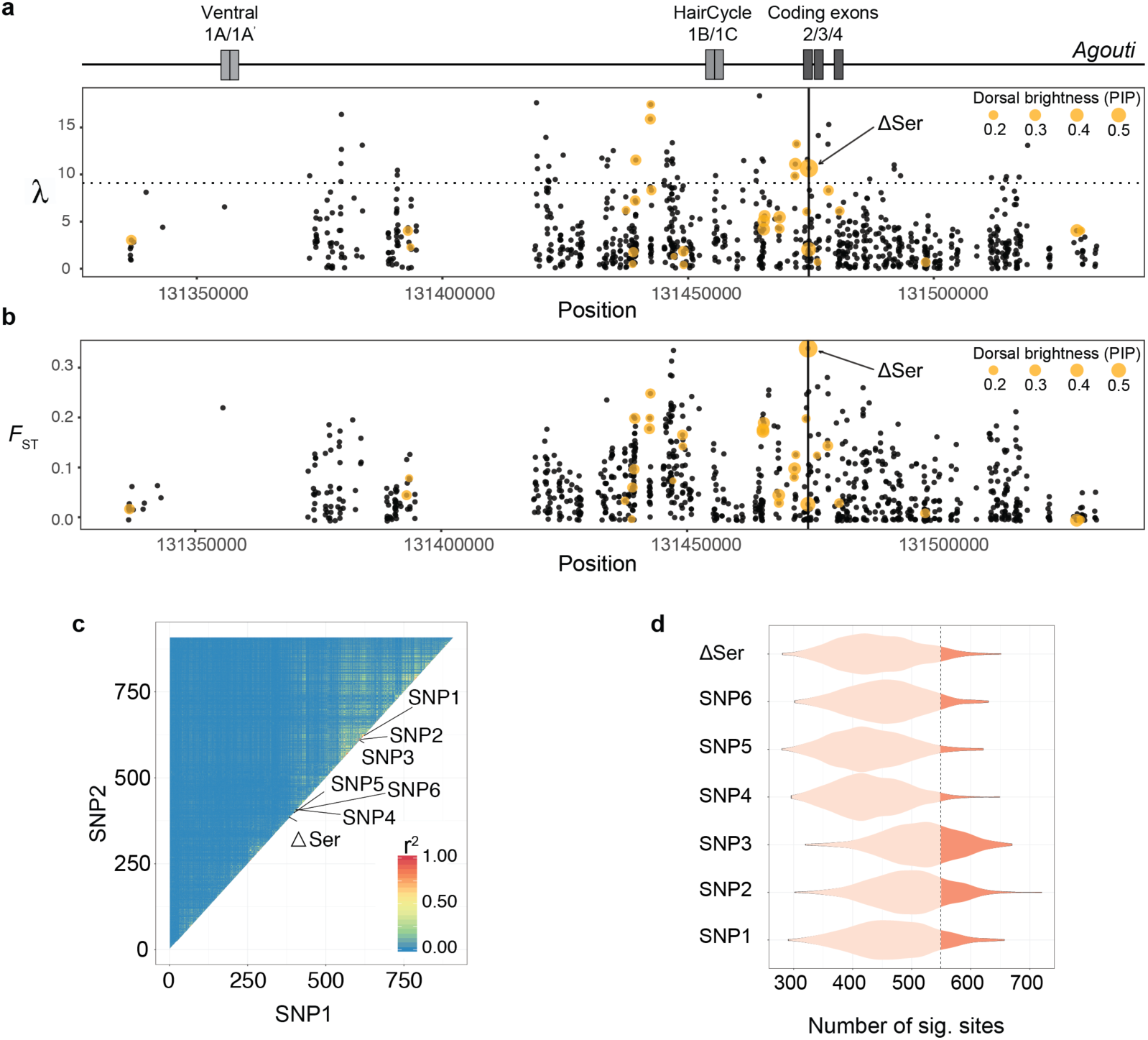
Candidate variants for selection within *Agouti*. **a**, Map of the *Agouti* locus showing non-coding exons (1A/1A’ responsible for ventral pigmentation; 1B/1C for banded dorsal hairs) and coding exons 2-4 (top). Likelihood ratio test statistic for identifying positive selection on *Agouti* in the dark soil enclosures (pooled data, bottom). Yellow dots indicate variants associated with dorsal brightness and their size indicates the relative strength of their associations (PIP). The dotted line represents the FDR-corrected threshold for all sites associated with dorsal brightness. Vertical line shows location of the ΔSer mutation. **b**, Variant-specific *F*_ST_ between populations from light and dark habitat used to colonize the dark soil enclosures. Variants associated with dorsal brightness are indicated as in panel (**a)**. **c,** LD heat map for all *Agouti* sites in pooled enclosures on dark soil. Sites with a minor allele frequency ≤ 10% were discarded. **d**, Expected number of significant sites at the 1% level when a single site is under selection. Distribution of the number of sites with a P value ≤ 0.01 when survivors are artificially resampled assuming a non-central sampling distribution with weights defined by the genotype at the target sites. Distributions are shown for the 7 candidate sites in the dark enclosures. The dashed vertical line indicates the observed number of sites with significant change. None of the 7 P values are significant after correcting for multiple testing (FDR).

To test whether selection on each of these candidate variants could account for the observed number of SNPs with biased genotype frequencies in the survivors, we recalculated null distributions by assigning each candidate individually as our single selected target site. We found that, after correction for multiple testing, each of the 7 could account for the observed change in genotype frequencies in the survivors (Fig. 4c). By contrast, a model using the SNP from the genome-wide control dataset with the most significant allele frequency change cannot explain the observed patterns (Extended Data Fig. 2b). Linkage disequilibrium analyses of the 7 candidate variants identified three linkage blocks (Extended Data Fig. 2c): two sets of three physically proximate regulatory SNPs and the ΔSer, the latter displaying low LD with all other candidate SNPs (Fig. 4d). These data suggest that each of these three linkage blocks harbor variants directly responding to selection on dorsal brightness. Thus, strong selection on a limited number of genetic targets within the *Agouti* locus is sufficient to drive extensive shifts in allele frequency and rapid change in phenotype.

## The functional and ecological effects of a deletion mutation in *Agouti*

To test the functional link between one of the variants in *Agouti* associated with survival and pigmentation, as well as uncover the causal molecular mechanism, we focused on the amino acid mutation ΔSer in *Agouti*. We chose this variant from of our 7 candidates because the ΔSer was highly associated with dorsal brightness (R^2^ = 0.11, *P* = 3.02e^−12^; Fig. 5a), shows a signature of selection in the enclosure populations (Fig. 4a) as well as in an admixed natural population^23^, and importantly, this mutation showed the highest level of genetic differentiation across the entire *Agouti* locus between mice that were originally captured from light and dark habitat (*F*_ST_ = 0.34; Fig. 4b; Extended Data Table 4). To determine whether the ΔSer alone has an effect on hair colour *in vivo*, we generated matching lines of transgenic lab mice (C57BL/6 mice, a strain with no endogenous *Agouti* expression) carrying the wildtype (WT) or the ΔSer *Peromyscus Agouti* cDNA, constitutively driven by the Hsp68 promoter (Fig. 5b). We used the ϕC31 integrase system, which produces single-copy integrants at the *H11P3* locus on mouse chromosome 11 to directly measure the effect of the *Agouti* ΔSer while avoiding variation caused by copy number, insertion site, or orientation of the construct^27^ (Extended Data Fig. 3a,b). Using a spectrophotometer to quantify differences in coat colour, we found that ΔSer mice had significantly lighter coats than mice carrying the wildtype *Peromyscus Agouti* cDNA (ΔSer vs WT, two-tailed *t*-test; *n* = 5, *P* = 0.001; Fig. 5c). Thus, the *Agouti* ΔSer mutation alone has a measurable effect on pigmentation and in the direction expected based on genotype-phenotype association data in natural *Peromyscus* populations.

**Figure 5.**
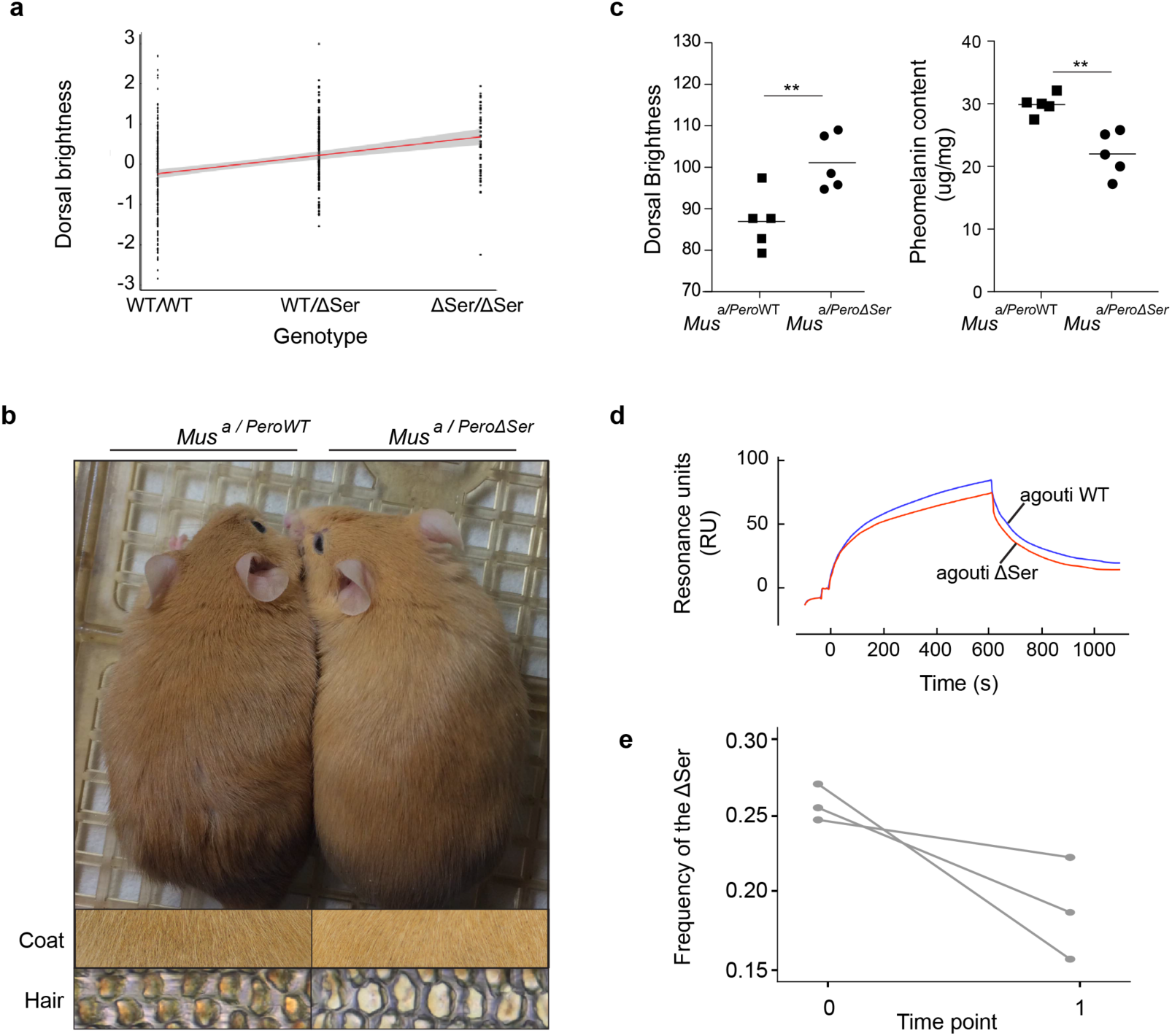
Phenotypic, molecular, and fitness effects of the Serine deletion. **a**, Linear regression of ΔSer genotypes and dorsal brightness; data pooled across all 6 enclosures. **b,** Matched lines of transgenic *Mus* (in C57BL/6, an *Agouti* knockout strain) expressing the wildtype (WT; dark) or the ΔSer (light) *Peromyscus Agouti* allele. Close-up pictures show the intensity of pheomelanin in dorsal coats and individual dorsal hairs from transgenic mice. **c,** Dorsal brightness (left) and benzothiazine-type pheomelanin degradation products (right) in the transgenic mice, measured with spectrophotometry and HPLC methods, respectively. **d,** Biochemical interaction of attractin and the N-terminal domain of the *Peromyscus* wildtype (blue) or the ΔSer (red) agouti protein. Values shown in arbitrary response units have been corrected for non-specific binding. **e**, Changes in *a*^*ΔSer*^ allele frequency across the 3 replicate dark enclosure populations.

To further characterize the phenotypic effects of the ΔSer variant, we examined and then quantified pigment in dorsal hair. Microscopic examination of individual hairs revealed that the hair of ΔSer mice contained a qualitatively lighter pigment than that of wildtype mice (Fig. 5b). To quantify this difference, we analysed pheomelanin content in the hair using chemical degradation products followed by high performance liquid chromatography^28-31^. We found that ΔSer mice had significantly lower amounts of pheomelanin (both benzothiazine- and benzothiazole-types) than hair from wildtype mice (ΔSer vs WT, two-tailed *t*-test; *n* = 5, *P* = 0.002; Fig. 5c and Extended Data Fig. 3c). These results indicate that the *Peromyscus* ΔSer causes a decrease in production of pheomelanin, which in turn causes hair to appear brighter overall.

The ΔSer is found in a highly conserved region of the N-terminal domain of the agouti protein, a region that directly binds to attractin, a transmembrane receptor expressed in melanocyte membranes and required for agouti function^32^. To understand the mechanism by which the ΔSer decreases pheomelanin production, we measured real-time binding interactions between the agouti and attractin proteins using Surface Plasmon Resonance (SPR). In SPR, one molecule (ligand) is immobilized on a sensor surface, while a potential interacting partner (analyte) is injected; reflection angle of polarized light from the sensor then serves as a proxy for the strength of the interaction between the molecules. For a ligand, we used the secreted isoform of natural human Attractin (ATRN^Ec^) and for the analyte, we used a synthetic version of the *Peromyscus* agouti wildtype or ΔSer N-terminal domain, a region known to retain full biochemical activity and bind attractin^32^. Application of wildtype or ΔSer agouti N-terminal domain to an attractin-coated chip produced sensorgrams characteristic of a biological interaction, approaching equilibrium over several minutes, and declining during washout to levels above baseline (Fig. 5d). However, we found that the wildtype N-terminal domain showed a stronger interaction with attractin relative to the ΔSer allele (Fig. 5d). We next estimated dissociation constants (K_d_), using Scatchard analysis of equilibrium binding levels at different concentrations, and showed that the wildtype domain has a nearly two-fold smaller K_d_ than the ΔSer domain (4.25 × 10^−7^ vs 6.94 × 10^−7^, respectively), consistent with the wildtype allele having a greater binding affinity to attractin (Extended Data Fig. 3d). Together, our genetic and biochemical experiments indicate that ΔSer causes lighter pigmented hair by decreasing the strength of the interactions with attractin, reducing pheomelanin production, and ultimately increasing the brightness of a mouse’s dorsal coat.

## Changes in the *Agouti* ΔSer allele through space and time

Based on its verified functional role in pigment variation, we next measured the frequency of the *Agouti* ΔSer allele across enclosures and over time. To both confirm the *Agouti* ΔSer genotype and to include individuals with missing data, we genotyped all individuals using a Taqman assay. The starting frequency of the ΔSer allele varied among the six enclosures, but on average was similar among dark and light enclosures (light enclosures mean = 29.85% ±1.80 S.E., dark enclosures mean = 25.79% ±0.68 S.E.). We observed idiosyncratic changes in allele frequency in the light enclosures, with two of three enclosures showing the expected increases in frequency of the ΔSer allele, but the degree of change was minor in all cases (average allele frequency change = 0.43% ±0.81 S.E.). By contrast, we observed significant decreases in the frequency of the ΔSer allele in all three replicate dark enclosures (average allele frequency change = 6.87% ±2.58 S.E.; Fig. 5e). This change in allele frequency amounts to a mean selection coefficient of 0.32 (±0.11 S.E.; one-sided t-test of selection coefficients in light versus dark enclosures: *t* = 2.990, df = 2.496, *P* = 0.037; Extended Data Table 5). Consistent with the negative phenotypic selection observed on light pigmentation in dark enclosures, these genetic results provide evidence for negative selection on the ΔSer allele associated with light pigmentation on dark substrates. Thus, by documenting allele frequency change over time, we demonstrate strong selection at the genetic level consistent with predictions based on the functional effects of the ΔSer variant.

## Discussion

Knowing the strength of selection in nature is essential for being able to predict rates of adaptive change^8,33-38^. We have a growing sense of the strength of selection acting on phenotypes^9,39-42^, and there is now extensive data providing statistical signatures of historical selection on the genome^43-46^. However, uncertainty remains concerning the magnitude and causes of genetic changes that occur as populations evolve under new ecological conditions^47-50^. Our experimental design mimics the replicated and reciprocal colonization of divergent habitats by populations carrying sequence variants that cause functional changes in a locally adapted phenotype. We have demonstrated that natural selection can result in rapid phenotypic change when appropriate standing genetic variation is available. Importantly, changes in both our focal trait (dorsal brightness) and components of its underlying genetic architecture (the ΔSer mutation) were highly predictable based on transgenic and biochemical assays as well as patterns of existing phenotypic and genotypic variation across habitat types. Together, these results suggest that knowledge about the functional connections between genotype, phenotype, and fitness could help predict future evolution under defined ecological conditions.

## ACKNOWLEDGEMENTS

We thank F. Baier, N. Bedford, A. Bendesky, J. Best, J. Chu, C. Clabaut, J. Gable, E. Hager, G. Hood, E. Jacobs-Palmer, E. Kay-Delaney, E. Kingsley, M. Manceau, N. Man in’t Veld, H. Metz, J-M. Lassance, E. Lievens, C. Linnen, N. Rubinstein, L. Schmitt, H. Wegener and I. Yen for field assistance; C. Clabaut, P. Muralidhar and K. Turner for laboratory assistance; Z. Gompert, B. Peterson, and J-M. Lassance for bioinformatics assistance; J. Demboski, B. Perrett, M. Perrett, B. Peterson, J. Ramos, L. Ramos, B. Ward, J. Wasserman, R. Wasserman, and the Denver Museum of Nature and Science for logistical support; and J. Chupasko for curatorial assistance. We thank G. Barsh, J. Losos, P. Nosil, D. Petrov, D. Schluter, and T. Thurman for commenting on the manuscript. R.D.H.B. was supported by a Natural Sciences and Engineering Research Council of Canada Banting Postdoctoral Fellowship and a Foundational Questions in Evolutionary Biology Postdoctoral Fellowship. S.L., M.F. and R.M., as well as laboratory work, were supported by a Swiss National Science Foundation Sinergia grant to J.D.J, H.E.H. and L. Excoffier. Fieldwork was funded by the National Geographic Society, Putnam Expedition Grants from the Harvard Museum of Comparative Zoology (MCZ), a Discovery Grant from the National Engineering and Research Council of Canada to R.D.H.B. H.E.H. is an Investigator of the Howard Hughes Medical Institute.

## AUTHOR CONTRIBUTIONS

R.D.H.B and H.E.H. designed and led the project. R.D.H.B. conducted the field experiment with C.C.Y.X. R.D.H.B conducted the phenotypic analyses and collected genomic data. S.P.P. and S.L. conducted the bioinformatics analyses. S.L., M.F. and J.D.J. conducted the statistical analyses. R.M. conducted the functional experiments, including the protein experiments with J.D.C and the melanin analysis with K.W. R.D.H.B. drafted the manuscript with major input from S.L., R.M., and H.E.H. All authors contributed revisions and approved the final version of the manuscript.

## Methods

### Enclosure construction and colonization

In May and June 2011, we constructed eight enclosures that contain 20,000 m^2^ of habitat: four on a light-substrate site on the Sand Hills (the ‘light site’; 42°35’13.44”N, 100°54’27.64”W) and four on a dark-substrate site off the Sand Hills (the ‘dark site’; 42°54’5.27”N, 100°30’32.30”W). Each enclosure is 25 m x 25 m with steel plate walls (1.5 m by 3 m plates of 24-guage galvanized steel) buried 0.75 m into the ground. Following a previous study^1^, we first trenched the perimeter of each enclosure to a depth of 0.75 m using a mechanical trencher equipped with a 15cm blade. We then buried steel posts (3.8 by 3.8 cm, 2.1 mm thick, and 2 m long) into the trench at 3 m intervals, allowing 1 m to protrude above ground. These posts served as supports for the walls, which we attached using self-tapping sheet metal screws. To extend the height of the walls to 1.8 m, steel posts (3.2 by 3.2 cm, 2.1 mm thick, cut to 1.2 m sections) were inserted 30 cm into the original support posts and secured using metal screws. We then attached chicken wire fencing (0.9 m high) to the extension posts with bailing wire to help exclude large mammals. Finally, we dug holes around the base of each post and filled with concrete for additional support. We removed native small mammals from each enclosure through repeated rounds of trapping using Sherman live traps.

Between June and August 2011, we sampled 481 mice using Sherman live traps from six locations within 50 km of each other (3 on light Sand Hills habitat and 3 on dark habitat, see Extended Data Table 6 for specific sampling locations; mice were collected under the Nebraska Game and Parks Commission Scientific and Educational Permit 901). We introduced between 75-100 of these mice to each of 6 of the 8 enclosures (3 at each site), with half coming from each habitat type. Prior to introduction, we anesthetized each mouse and recorded sex, age, length, weight, and pigmentation phenotype (see below). We did not detect significant differences in any non-pigment related traits between mice captured on light versus dark habitat. To allow for individual identification, we marked the ears and injected a small radio ID-tag between the scapula of each mouse. To collect DNA, we took a 1-mm diameter tail tip, which was preserved in ethanol.

### Coat colour variation in experimental populations

To measure pigmentation among the experimental populations of deer mice, we used a spectrophotometer to measure reflectance of each mouse at 9 locations across the body (3 measurements along the dorsum, flank, and ventrum), as described previously^2^. We then used Color Analysis Programs v1.05^3^ to process the raw SpectraSuite files and extracted 7 colour summary statistics. We computed averages for each of these summary statistics for each of the body regions (dorsum, flank, ventrum), resulting in a total of 21 colour variables for each mouse. After normal quartile transformation on each variable, we performed principle components analysis (PCA). After examining the scree plot, we selected the first 5 principal components (PCs), which together accounted for 84% of the variation in the data. Because factor loadings were not easily interpretable in the unrotated data, we performed a varimax rotation on the first 5 PCs^4^. Based on their loadings, the top 5 PCs corresponded to dorsal/flank brightness and contrast, dorsal/flank hue, ventral chroma and hue, ventral brightness and contrast, and dorsal/flank chroma and hue. For simplicity, we refer to these PCs in the main text as dorsal brightness, dorsal hue, ventral chroma, ventral brightness, and dorsal chroma. Higher values of each trait correspond to lighter phenotypes.

### Colour pattern variation in experimental populations

We measured two aspects of colour pattern: dorsal-ventral boundary and tail pattern. We took digital images of each mouse using a Canon EOS Rebel T3i (Canon U.S.A., Lake Success, NY) with a Sigma 18-250mm lens (Sigma U.S.A., Ronkonkoma, NY) and a X-Rite ColorChecker Passport (X-Rite, Grand Rapids, MI). We imported tiff files into ImageJ and outlined the dorsum (coloured portions of the dorsum, with legs and tail excluded) and body (outlined entire mouse, legs and tail excluded). Dorsal-ventral boundary was calculated as (body-dorsum)/body, followed by a normal quartile transformation. We used the following qualitative scores for tail pattern: 0 = no coloured portion, 1 = basal portion of tail is coloured, 2 = coloured portion extends to approximately half the length of the tail, 3 = intermittent colour throughout full length of tail, 4 = full colouration of tail, following previous work^5^.

### Population sampling in the enclosures over time

Following the introduction of mice to the enclosures, we conducted 5 sampling trips (August 2011, October 2011, December 2011, April 2012, August 2012) to trap mice in the enclosures and determine survivorship of individuals in each population (individual identities were verified by use of implanted radio tags and a handheld reader). We used 50 traps per enclosure and conducted up to 40 trapping sessions in each enclosure each trip. Using Huggin’s closed population capture-recapture model^6,7^ in R package mra version 2.16.11, our recapture rates ranged from 90 - 99% (average = 95%).

### Measuring natural selection on pigmentation traits

We used selection analyses^8-10^ to characterize the form, intensity, and direction of selection acting on all traits we had *a priori* reason to suspect as being important to survival^2,11^ - that is, those traits found to be significantly different between mice captured on light versus dark habitat: dorsal brightness, dorsal chroma, ventral brightness, ventral chroma, and tail pattern. For all phenotypic analyses, we used only adult mice as coat colour and pattern can change from juvenile to adult pelage, and colour variation is low among juvenile mice (most juveniles are a consistent grey colour). However, including juveniles in the analyses did not change the direction of selection for any trait.

For each enclosure, we estimated the fitness function relating survival (0 or 1) to the five traits using a generalized additive model (GAM) and the default settings in the function gam in the R package MGCV version 1.8-12^12^. We then used the function gam.gradients in the R package GSG version 2^10^ to obtain directional and quadratic selection gradients. We evaluated the statistical uncertainty in the estimated selection gradients using 1000 bootstrap replicates. Following the first time period, there was sufficient mortality (Fig. 2) that population sizes were not large enough to fit a GAM to all phenotypic traits (there were more coefficients in the model than data), and thus we did not continue our selection analyses past the first re-sampling point.

The methods above identified dorsal brightness as the only trait with consistent evidence for selection across multiple enclosures. To visualize selection on this trait without *a priori* assumptions about the shape of the fitness function, we produced nonparametric fitness surfaces for each treatment^9,13^. We generated predicted values and Bayesian standard errors for the cubic splines using a GAM in the function gam of the R package MGCV version 1.8-12^12^. Smoothing parameters were chosen from values that minimized the generalized cross-validation (GCV) scores. We then visualized fitness surfaces by plotting predicted values with 1 Bayesian standard error, using the inverse of the logit link function to obtain the original scale.

### DNA sequencing library preparation

To prepare sequencing libraries, we used DNeasy kits (Qiagen, Germantown, MD) to extract DNA from tail samples. We then prepared genomic libraries following the SureSelect Target Enrichment Protocol v.1.0, with some modifications. In brief, 10-20 µg of each DNA sample was sheared to an average size of 200 base pairs (bp) using a Covaris ultrasonicator (Covaris Inc., Woburn, MA). Sheared DNA samples were purified with Agencourt AMPure XP beads (Beckman Coulter, Indianapolis, IN) and quantified using a Quant-iT dsDNA Assay Kit (ThermoFisher Scientific, Waltham, MA). We then performed end-repair and adenylation, using 50 µl ligations with Quick Ligase (New England Biolabs, Ipswich, MA) and paired-end adapter oligonucleotides manufactured by Integrated DNA Technologies (Coralville, IA). Each sample was assigned one of 48 5-bp barcodes. We pooled samples into equimolar sets of 9-12 and conducted size selection of a 280-bp band (+/-50-bp) on a Pippin Prep with a 2% agarose gel cassette. We performed 12 cycles of PCR with Phusion High-Fidelity DNA Polymerase (NEB), following manufacturer guidelines. To permit additional multiplexing beyond that permitted by 48 barcodes, we also added one of 12 6-bp indices to each pool of 12 barcoded samples. Following amplification and AMPure purification, we assessed the quality and quantity of each pool (17 total; see below) with an Agilent 2200 TapeStation (Agilent Technologies, Santa Clara, CA) and a Qubit-BR dsDNA Assay Kit (Thermo Fisher Scientific, Waltham, MA).

### DNA enrichment and sequencing

To enrich sample libraries for both the *Agouti* locus as well as randomly distributed genomic regions, we used a MYcroarray MYbaits capture array (MYcroarray, Ann Arbor, MI). In brief, this probe set includes the 185-kb region containing all known *Agouti* coding exons and regulatory elements (based on a *P. maniculatus Agouti* BAC clone^14^) and >1000 non-repetitive regions averaging 1.5-kb in length at random locations across the *P. maniculatus* genome. We enriched for regions of interest following the standard MYbaits v.2 protocol for hybridization, capture, and recovery. We then performed 14 cycles of PCR with Phusion High-Fidelity Polymerase and a final AMPure purification. We assessed final quantity and quality using a Qubit-HS dsDNA Assay Kit and Agilent 2200 TapeStation.

After enriching our libraries for regions of interest, we combined them into 17 pools and sequenced across 10 lanes of 125-bp paired-end reads using an Illumina HiSeq2500 platform (Illumina, San Diego, CA). From the obtained read pairs, 94% could be confidently assigned to individual samples (i.e., excluding reads when either barcodes or indices were ambiguous and/or had low base qualities), resulting in an average read pair count of 2,674,111 per individual.

### Sequence alignment

Following our previous approach^15^, we first removed contaminants and low-quality nucleotides. Partially error-corrected, single-end reads were generated from each read pair using FLASH v.1.2.11^16^ and aligned to the repeat-masked *Peromyscus maniculatus bairdii* Pman_1.0 reference assembly (http://www.ncbi.nlm.nih.gov/assembly/GCF_000500345.1/). More than 98% of all reads mapped to the reference genome. Finally, we removed duplicates, conducted multiple sequence alignments, and adjusted base qualities.

### Variant Calling and Filtering

Following^15^, variants were called per individual, and diploid genotypes were jointly inferred per enclosure pair (*i.e.*, one ‘light’ and one ‘dark’ enclosure with n_1_=203 individuals, and n_2_=147, n_3_=122). Post-genotyping, we excluded 10 individuals with more than 80% missing data and merged all enclosures into a pooled dataset. A principal component analysis (PCA) of the data revealed the presence of 16 outlier individuals that were then also excluded. We further restricted this pooled dataset to biallelic variants that were genotyped in at least 80% of individuals. In the absence of a validation dataset for this species, we applied stringent hard filter criteria to avoid false positives due to sequencing errors or alignment issues (see^15^ for details). The final call set contained 2,442 variants for the *Agouti* scaffold and 53,507 for the genome-wide control dataset. We conducted analyses of single enclosures by sub-setting the pooled data with the appropriate list of individuals, such that the same set of variants could be compared across all soil colours and replicates.

### Standardizing the SNP dataset across enclosures

For each treatment (soil colour), we created a pooled dataset consisting of all SNPs present in the 3 enclosures. We applied a missing data filter to both pooled datasets independently. For each treatment, we removed individuals with more than 80% missing nucleotide sites. Among the remaining individuals, we removed sites with more than 20% missing data. Finally, we conducted enclosure-specific analyses by back-partitioning the pooled data such that the same set of SNPs could be compared among enclosures.

### Calculation of probabilities of genotype distributions in surviving individuals

To test the null hypothesis that survival and genotype were independent, we calculated for every variant the probability that genotype frequencies in the survivors were the result of unbiased (neutral) versus biased (selective) sampling using the multivariate Wallenius non-central hypergeometric distribution^17^. From this analysis, we obtained the number of sites with a P value < 0.01 assuming unbiased sampling. To test whether this number was a consequence of conducting a large number of statistical tests, we next calculated the distribution of the number of sites with a P value < 0.01 expected under neutrality. Finally, to test whether the number of significant sites could be inflated due to linkage, we calculated the null distribution of sites with a P value < 0.01 assuming a model with a single site under selection. A detailed description of this statistical approach is provided in the Supplemental Methods. To test for significant selection at each locus, we calculated the ratio of biased over unbiased probabilities. We obtained P values for this probability ratio using a chi-square distribution with 2 degrees of freedom and set the significance threshold to 0.05 after FDR correction. The R code implementing these analyses is available from (https://gitlab.mpcdf.mpg.de/laurent/survseq). We calculated *F*_ST_ and LD using vcftools (–hap-r2 option, version 0.1.15). The code for the visualization in Figure 4d is modified from https://stackoverflow.com/questions/22278951/combining-violin-plot-with-box-plot.

### Functional testing of the *Agouti* mutation using site-specific transgenesis

To explicitly test if the Serine deletion mutation (ΔSer) in *Agouti* exon 2 has a measurable effect on pigmentation, we used a recombinase mediated cassette exchange approach to insert the *Peromyscus* wildtype (WT) and ΔSer alleles into the *Mus* C57BL/6, a strain that contains a knockout *Agouti* mutation.

#### Plasmids

To construct site-specific insertion plasmids, we synthesized a 1.1-kb fragment corresponding to the Hsp68 promoter and the *Peromyscus maniculatus Agouti* cDNA wildtype (WT) sequence by GenScript and then cloned these fragments into the PUC19 vector. From the WT template sequence, we deleted the Serine amino acid using a site-directed mutagenesis kit (New England Biolabs) to generate the ΔSer construct. Thus, the WT and ΔSer constructs are identical, except for the deletion of a single amino acid Serine at amino acid position 48. We digested the resulting vectors with ClaI and HindIII and then cloned these fragments into the pBT378.2 plasmid (Applied Stem Cell), which contains two AttB sites. We confirmed the plasmid sequences by Sanger sequencing.

#### Site-specific integration into H11P3 sites

The *Agouti* WT and ΔSer circular plasmids were individually mixed with *in vitro* transcribed φC31 integrase mRNA and microinjected by Applied Stem Cell into the pronuclei of heterozygous H11P3 embryos, harvested from C57BL/6 (B6) mice. In the presence of φC31 integrase, the attB sequences from the donor vector can undergo site-specific recombination with the attP sequences in the H11P3 mouse locus^18^. We transferred injected zygotes to the oviduct of CD1 foster mice and identified founder animals by PCR-based genotyping with primer pairs PR425N/*Agouti*-WT.v2-5GoI-R; *Agouti*-WT.v2-F2/*Agouti*-WT.v2-R2; *Agouti*-WT.v2-3GoI-F/Frt-R1 to screen for site-specific and random integrations (see Extended Data Figure 3, Extended Data Table 5).

We then crossed founders containing site-specific insertions in the same orientation each to C57BL/6J mice to produce offspring in which we could characterize pigmentation phenotypes (as described above) and quantify the amount of pigment in hairs (see below).

### Quantifying pheomelanin production

To measure the amount of phaeomelanin in hairs, we homogenized hair samples (ca. 15 mg) with a Ten-Broeck glass homogenizer at a concentration of 10 mg/mL. We then subjected aliquots of 100 μL to hydroiodic acid hydrolysis^19^ and alkaline hydrogen peroxide oxidation^20^. We analysed benzothiazine-type pheomelanin (BT-PM) as a specific degradation product of 4-amino-3-hydroxyphenylalanine (4-AHP) produced by the hydroiodic acid hydrolysis, while benzothiazole-type pheomelanin (BZ-PM) was quantified as thiazole-2,4,5-tricarboxylic acid (TTCA), produced by the alkaline hydrogen peroxide oxidation. Finally, we calculated the BT-PM and BZ-PM content by multiplying the 4-AHP and TTCA contents by factors of 7 and 34 respectively^20-22^.

### Measuring interactions between agouti and attractin using surface plasmon resonanc

We used surface plasmon resonance (SPR) analysis on a BIAcore 3000 instrument to test for interactions between the agouti N-terminal WT and N-terminal ΔSer peptides with attractin ectodomain. Flow cells within a CM5 biosensor chip (GE Healthcare Life Sciences) were activated with N-hydroxysuccinimide (NHS) in the presence of 1-ethyl-3-(3-dimethylaminopropyl) carbodiimide hydrochloride generating an NHS ester that bound to free amines on full-length recombinant human Attractin ectodomain^23^ passed over the surface (25 µg/ml in 100mM sodium acetate, pH 4.5). We blocked free NHS ester on the flowcell surface using ethanolamine (1 M). We next exposed an additional cell to immobilizing and blocking reagents in the absence of protein as a control surface. After extensive washing of the surfaces with binding buffer (10 mM HEPES, 150 mM NaCl, pH 7.0, 0.05% Tween 20), we assessed binding of the respective agouti peptides by injecting varying concentrations (0-312.5nM) simultaneously over the control and attractin surfaces. Note that the ectodomains of mouse and human Attractin share 94% identity, and full-length mouse agouti binds to human Attractin ectodomain with an apparent K_D_ of < 50nM. Between each cycle, we regenerated the surface using 20 mM glycine-HCl, pH 2.8. We corrected values shown in Resonance Units (RU) for non-specific binding by subtracting the SPR of the control flow chamber exposed to the same injected material followed by subtraction of buffer alone passing over the attractin surface. We analysed the resulting data using BIAevaluation 3.2 software and fitted the data to a 1:1 Langmuir binding model with separate k_d_ and k_a_ determinations. Finally, we calculated the dissociation constant (K_D_) as k_d_/k_a_, and confirmed by manual linear transformation of the binding isotherms.

### Genotyping the Serine deletion

To genotype all founding individuals for the ΔSer mutation, we designed a Custom TaqMan genotyping assay targeting WT and ΔSer *Agouti* sequences^2^, using the TaqMan Custom Assay Design Tool (Thermo Fisher Scientific, Waltham, USA). We then used the resulting primers and probes with 2X TaqMan Genotyping Master Mix (Thermo Fisher Scientific), following the manufacturer’s protocol, to amplify and genotype samples in 96-well plates on a Mastercycler® RealPlex2 (Eppendorf, Hamburg, Germany).

### Estimating selection on Serine deletion alleles and genotypes

To estimate the selection coefficient (*s*) for viability selection on the WT and ΔSer alleles in each replicate enclosure population, we calculated the change in frequency of each allele relative to the change in frequency of the WT allele, subtracted from 1, following^24^. Similarly, *s* for genotypes was calculated as the change in frequency of the homozygous ΔSer genotype relative to the change in frequency of the homozygous WT genotype minus 1, when selection favored the homozygous ΔSer genotype. When there was evidence for selection against the homozygous ΔSer genotype, we calculated *s* as the change in frequency of the homozygous ΔSer genotype relative to the change in frequency of the homozygous WT genotype, subtracted from 1. This produces selection coefficients for the homozygous ΔSer genotype when it is favored and against the homozygous ΔSer genotype when it is selected against, using the homozygous WT genotype as the reference. We calculated *h* (the dominance coefficient) as the change in frequency of the heterozygous genotype subtracted from 1 divided by *s* of the homozygous ΔSer genotype^24^. Standard errors for *s* and *h* are from measurements of *n* = 3 dark enclosures.

### Data and code availability

Data supporting the findings of this study are available in the manuscript, its supplementary files, and from the corresponding authors on request. We have deposited sampling, phenotype, and survival data in the Dryad Digital Repository (DOI: XXXXXXX) and sequence data in the NCBI Short Read Archive with the primary accession code SUB4114786. The R code implementing analyses of genotype distributions is available from https://gitlab.mpcdf.mpg.de/laurent/survseq.

## Extended Data Tables

**Extended Data Table 1.** Pair-wise correlation between pigment traits.

| Trait 1 | Trait 2 | Slope | $r^2$ | P value |
| --- | --- | --- | --- | --- |
| Dorsal brightness | Dorsal chroma | -0.028 | 0.059 | 0.632 |
| Dorsal brightness | Ventral brightness | 0.030 | 0.054 | 0.576 |
| Dorsal brightness | Ventral chroma | -0.015 | 0.053 | 0.779 |
| Dorsal brightness | Tail pattern | -0.074 | 0.050 | 0.142 |
| Dorsal chroma | Ventral brightness | 0.002 | 0.050 | 0.965 |
| Dorsal chroma | Ventral chroma | 0.111 | 0.049 | 0.024* |
| Dorsal chroma | Tail pattern | -0.118 | 0.046 | 0.010* |
| Ventral brightness | Ventral chroma | 0.055 | 0.054 | 0.307 |
| Ventral brightness | Tail pattern | -0.108 | 0.051 | 0.034* |
| Ventral chroma | Tail pattern | -0.074 | 0.051 | 0.153 |

\* = $P < 0.05$ , F-test

**Extended Data Table 2.** Standardized quadratic selection gradients (ϒ) for pigmentation traits.

| Trait | Dark site |  |  | Light site |  |  |
| --- | --- | --- | --- | --- | --- | --- |
|  | Enclosure<br>1 | Enclosure<br>2 | Enclosure<br>3 | Enclosure<br>1 | Enclosure<br>2 | Enclosure<br>3 |
| Dorsal<br>brightness | -0.037<br>(0.534) | 0.021<br>(0.051) | -0.005<br>(0.077) | 1.013<br>(3.985) | 0.202<br>(0.162) | 0.143<br>(0.467) |
| Dorsal<br>chroma | -0.006<br>(0.065) | 0.008<br>(0.022) | 0.000<br>(0.010) | 0.037<br>(0.727) | 0.002<br>(0.047) | 0.047<br>(0.269) |
| Ventral<br>brightness | -0.019<br>(0.748) | 0.000<br>(0.007) | -0.001<br>(0.013) | 0.050<br>(0.642) | 0.005<br>(0.038) | 0.144<br>(1.297) |
| Ventral<br>chroma | -0.011<br>(0.413) | 0.001<br>(0.014) | -0.003<br>(0.034) | 0.000<br>(0.754) | 0.025<br>(0.052) | 0.008<br>(0.116) |
| Tail<br>pattern | 0.000<br>(0.574) | 0.002<br>(0.014) | 0.000<br>(0.017) | 0.016<br>(199.713) | 0.057<br>(0.097) | 0.067<br>(0.436) |

Parentheses give 1 standard error. Standard errors and P values determined by 1000 bootstrap replicates. All selection gradients calculated over the first three months following introduction to the enclosures. $P > 0.05$ in all cases.

**Extended Data Table 3.**
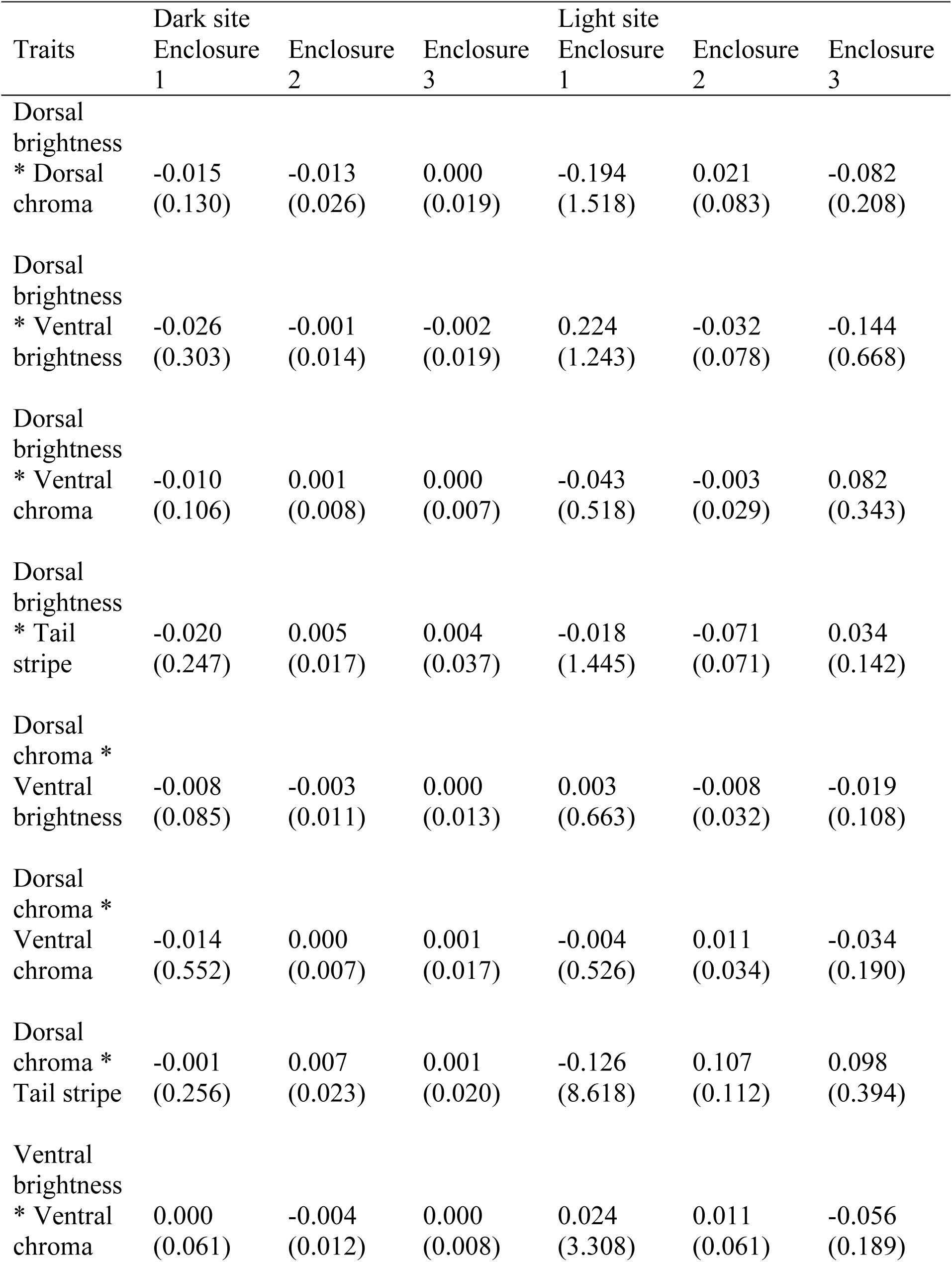
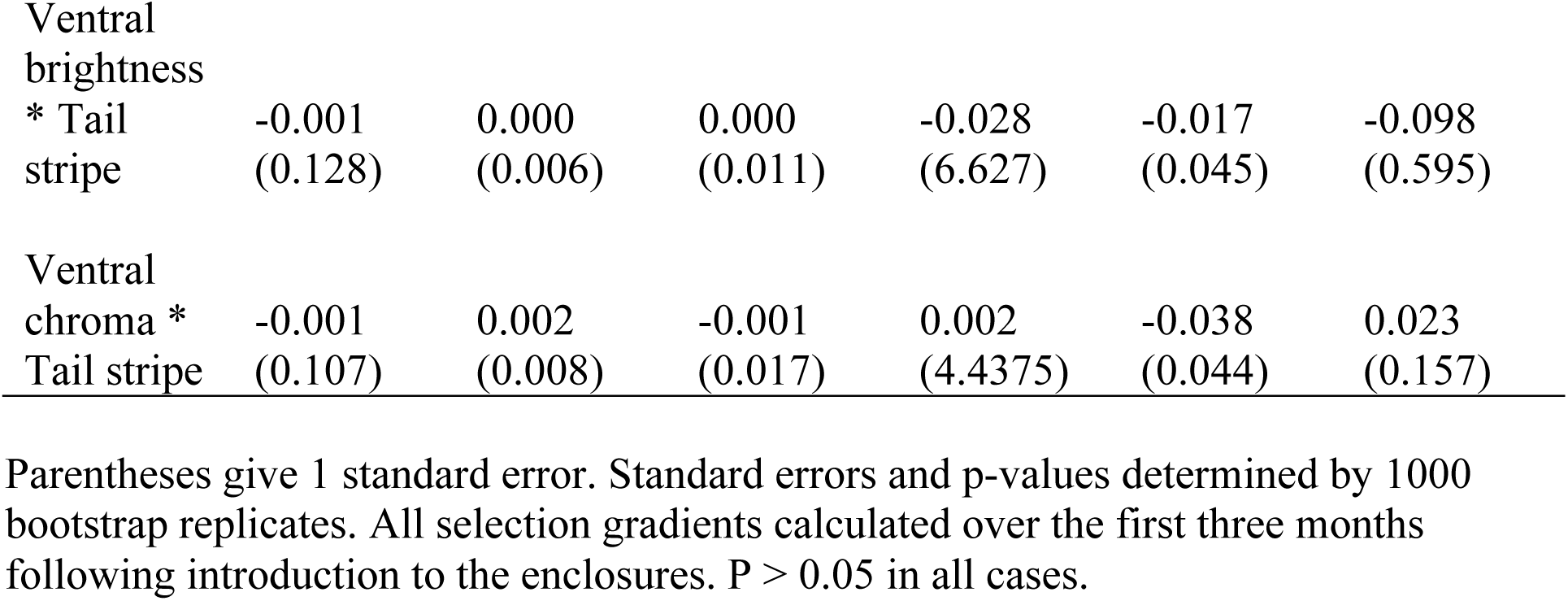
Standardized quadratic selection gradients (ϒ) for correlational selection on pigmentation traits.

**Extended Data Table 4.** Candidate SNPs for targets of direct selection.

| Variant | Chromosome | Position | Initial frequency | Frequency change<sup>1</sup> | PIP<sup>2</sup> | LRT | P value<sup>3</sup> | P value<sup>4</sup> | $F_{st}$<sup>5</sup> |
| --- | --- | --- | --- | --- | --- | --- | --- | --- | --- |
| SNP1 | chr4 | 131439350 | 0.43 | -0.08 | 0.16 | 11.54 | 0.02 | 0.11 | 0.20 |
| SNP2 | chr4 | 131442316 | 0.39 | -0.09 | 0.16 | 15.89 | 0.01 | 0.06 | 0.18 |
| SNP3 | chr4 | 131442394 | 0.39 | -0.09 | 0.11 | 17.43 | 0.01 | 0.05 | 0.20 |
| SNP4 | chr4 | 131471739 | 0.28 | -0.06 | 0.12 | 9.84 | 0.03 | 0.14 | 0.08 |
| SNP5 | chr4 | 131471821 | 0.31 | -0.07 | 0.21 | 11.11 | 0.02 | 0.11 | 0.10 |
| SNP6 | chr4 | 131472069 | 0.31 | -0.08 | 0.12 | 13.25 | 0.01 | 0.10 | 0.13 |
| ΔSer | chr4 | 131474564 | 0.26 | -0.06 | 0.54 | 10.67 | 0.02 | 0.12 | 0.34 |

1: Mean allele frequency change in the dark enclosures
2: Posterior inclusion probability (PIP) indicating the strength of the association with dorsal brightness
3: FDR-corrected P value of the LRT test when considering only variants in *Agouti* with a MAF >10% and associated with dorsal brightness
4: FDR-corrected P value of the LRT test when considering all variants in *Agouti* with a MAF >10%
5: Weir and Cockerham $F_{st}$ between mice captured in light versus dark habitat

**Extended Data Table 5.**
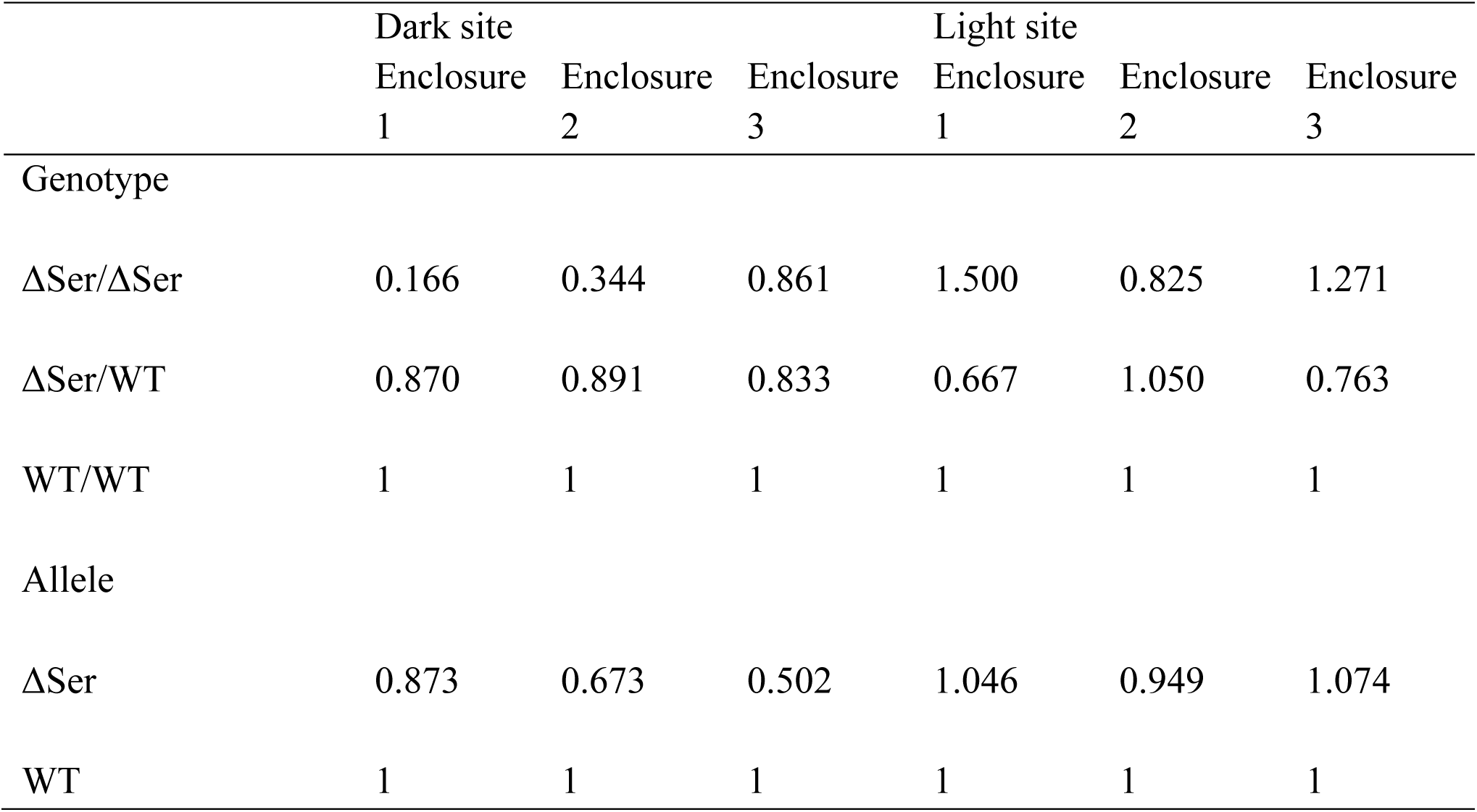
Relative fitness of Serine deletion genotypes and alleles.

**Extended Data Table 6.**
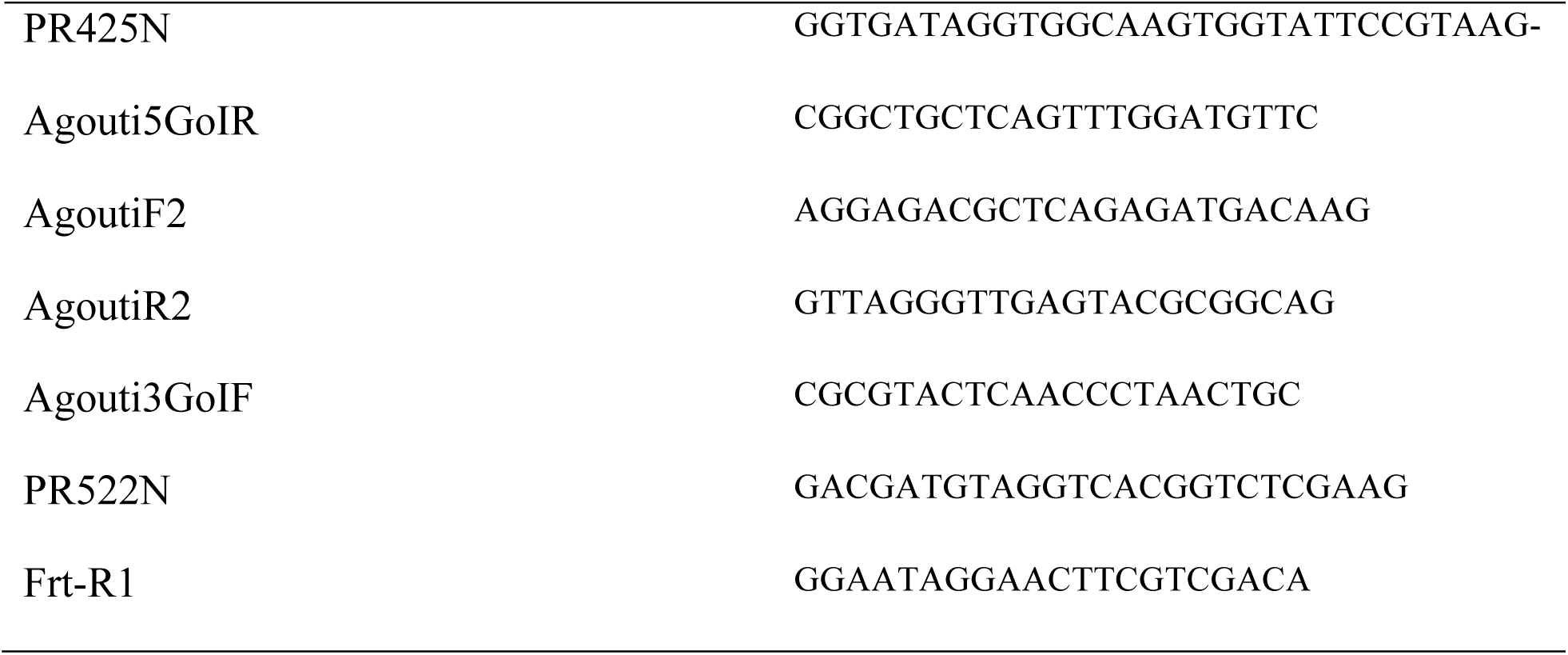
Primers used for genotyping the transgenic *Mus* lines.

**Extended Data Table 7.** Sampling locations of mice.

| Location | Full name | Habitat type | Coordinates |
| --- | --- | --- | --- |
| MER | Merritt Reservoir | Light | 42 35'42" 105 55'09" |
| BILL | Bill's field | Dark | 42 54'05" 100 30'32" |
| DAN | Danielski's field | Dark | 42 50'48" 100 34'01" |
| SCH | Schlagel creek | Light | 42 42'07" 100 37'00" |
| SMF | Samuel McKelvie National Forest | Light | 42 39'09" 100 56'37" |
| LON | Lonnie's field | Dark | 42 54'34" 100 34'37" |

**Extended Data Figure 1.**
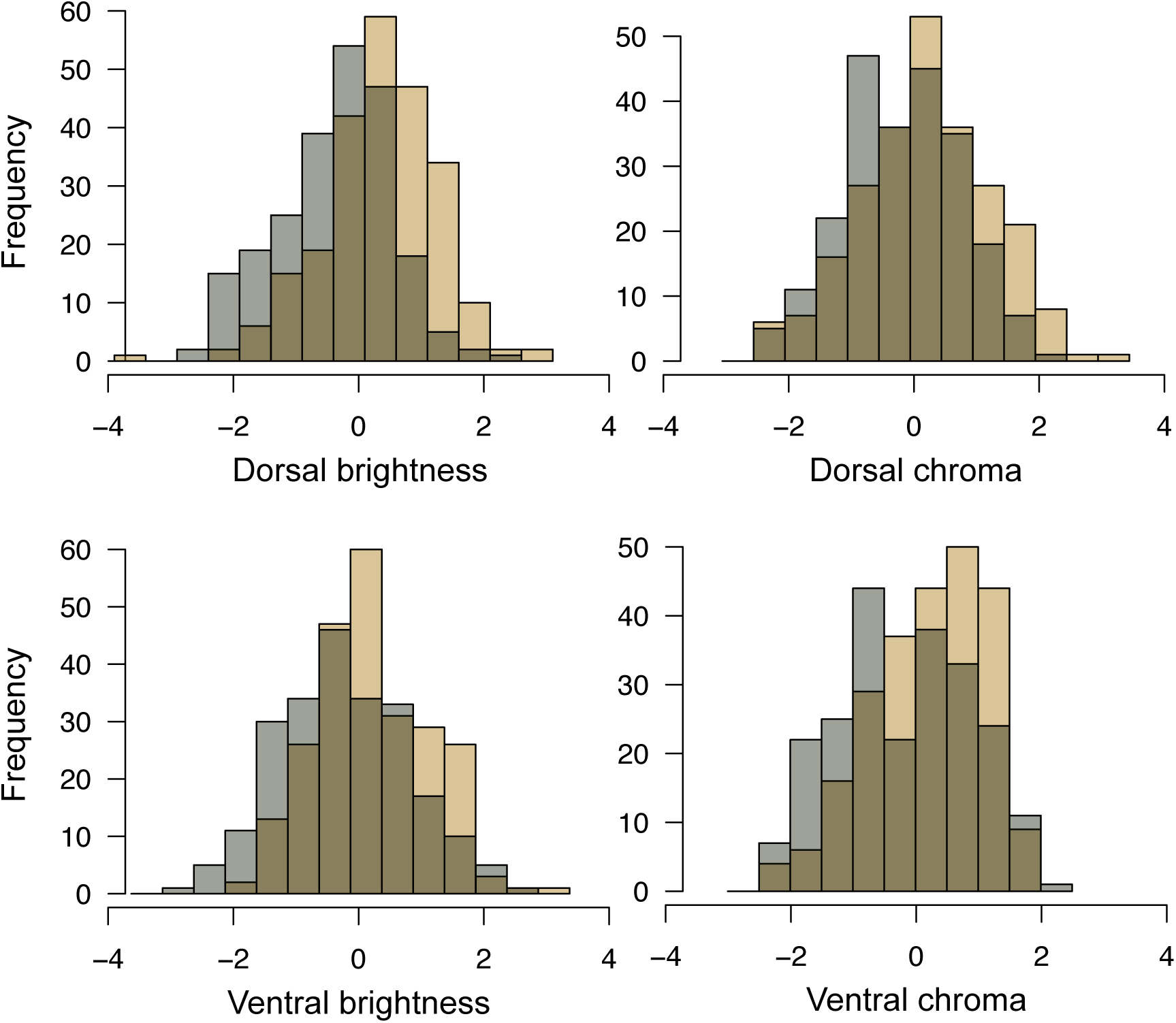
Distributions of quantitative pigmentation traits for mice captured on light (tan) versus dark (grey) habitat. Higher scores reflect lighter pigmentation (see Methods). All traits showed significant differences between mice captured on light versus dark habitat (two-sided t-tests, P <0.001).

**Extended Data Figure 2.**
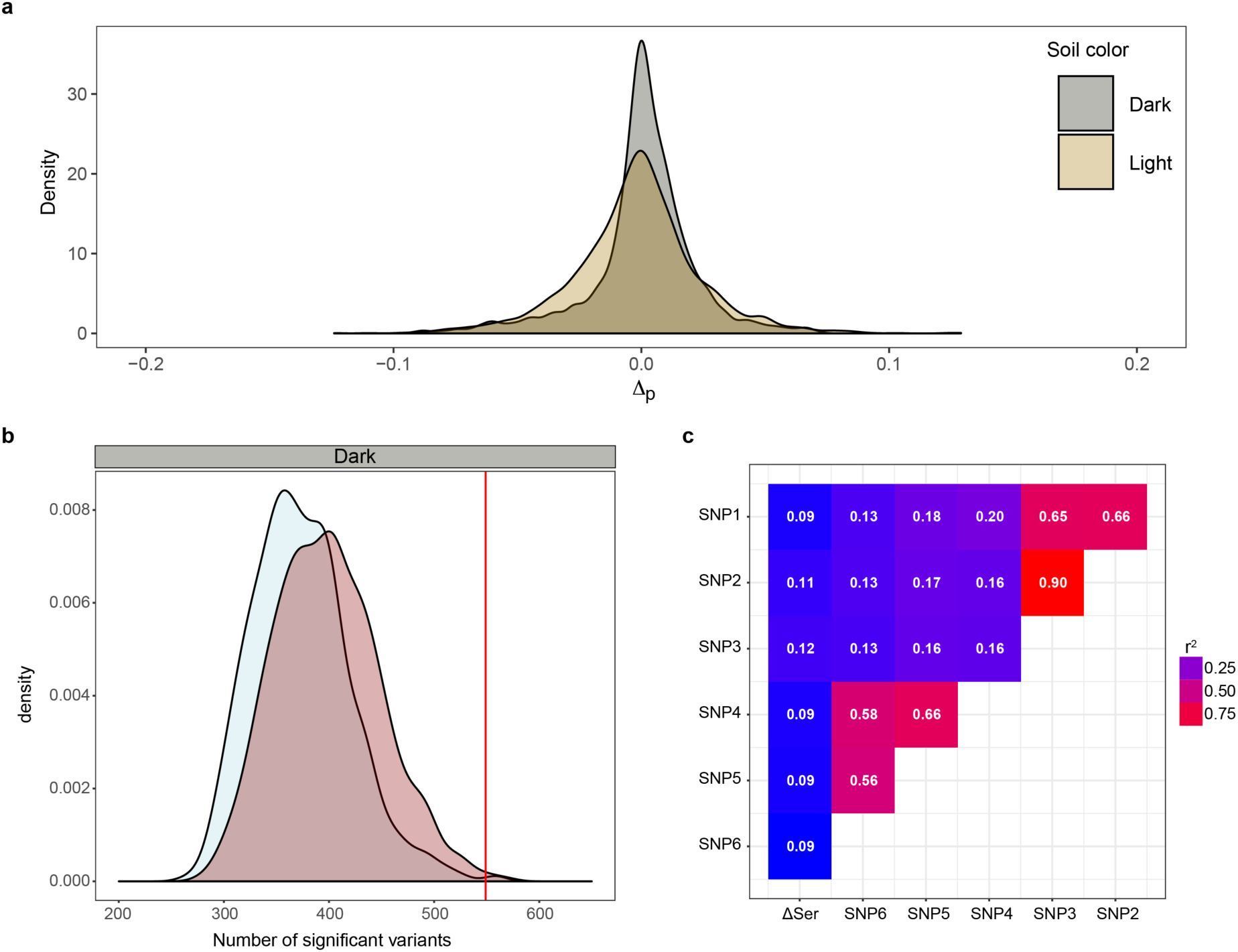
Allele frequency change and relationship between variants in *Agouti*. **a**, Distribution of allele frequency changes at the *Agouti* locus in the light (tan) and dark (grey) enclosures (pooled). The larger frequency changes in light enclosures are consistent with the lower number of survivors and therefore higher sampling error. **b**, Null distributions of the number of sites in *Agouti* expected to show significant allele frequency change at the 1% level in the dark enclosures. For the selection model, the site under selection is the site in the genome-wide control data with the lowest probability of having genotype frequencies in the survivors that are the result of random sampling. The observed number of significant sites is indicated by the vertical line. Note truncated x-axis. **c**, Linkage disequilibrium (r^2^) between the 7 candidate *Agouti* variants for selection listed in Extended Data Table 4.

**Extended Data Figure 3.**
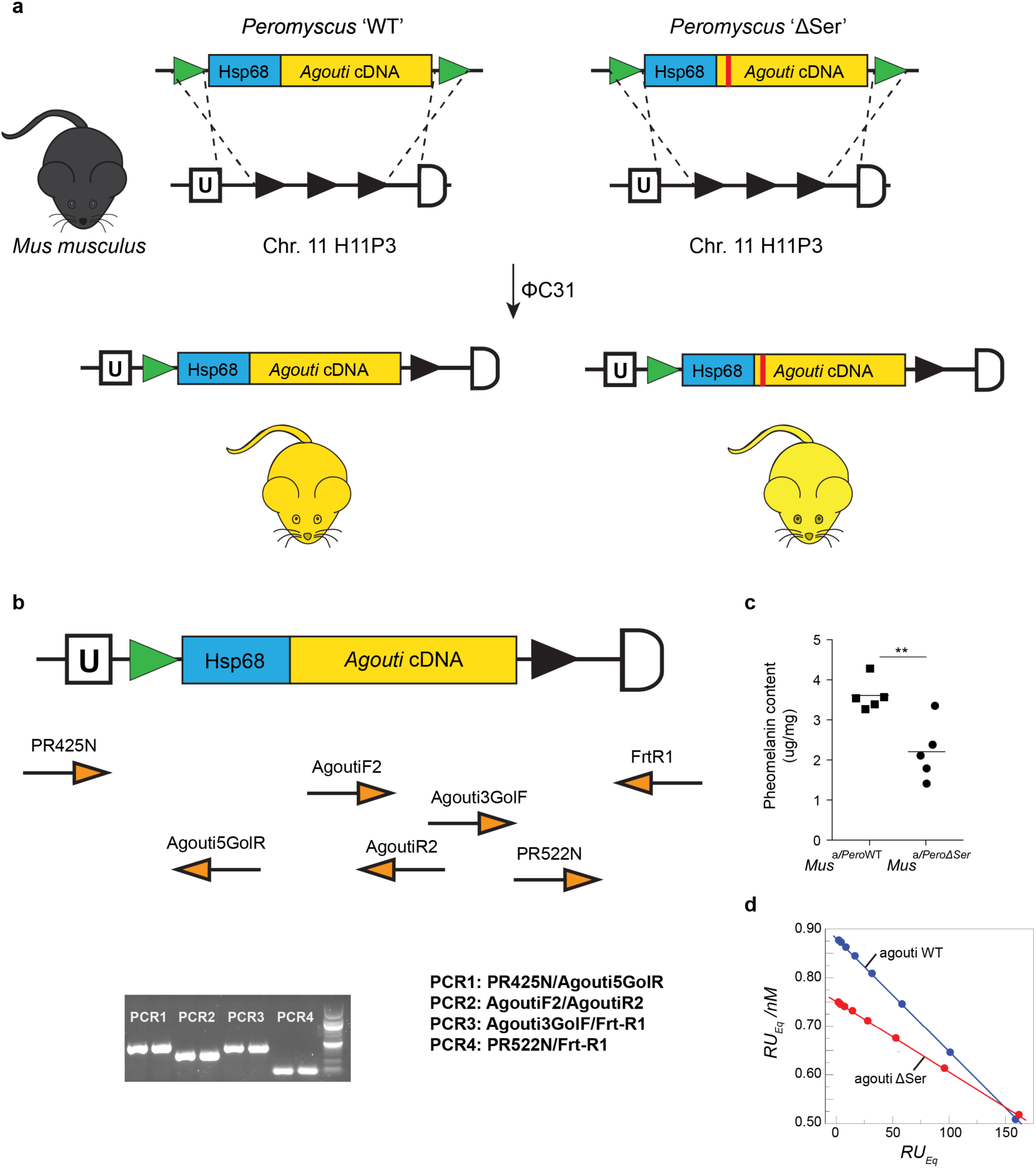
Functional experiments. **a,** In the presence of φC31 integrase, plasmids containing the Hsp68 promoter and the *Peromyscus Agouti* wildtype or ΔSer cDNA, as well as AttB, undergo site-specific recombination with the attP sequences of the H11P3 mouse locus and integrate into the genome. Founders have a mosaic black and yellow pattern but subsequent generations constitutively express agouti and have yellow coats. **b,** Genotyping strategy used. See Extended Data Table 5 for primer sequences. **c,** Pheomelanin degradation products (benzothiazole-type) in the transgenic mice, measured with spectrophotometry and HPLC methods (ΔSer vs WT, two-tailed *t*-test; *n* = 5, *P* = 0.006). **d,** Scatchard linear analysis was used to perform equilibrium binding analysis and determine the dissociation constant (K_d_) for the interactions between the N-terminal domain of the *Peromyscus* wildtype (blue) or the ΔSer (red) agouti protein.

## Application of the Wallenius’ non-central hypergeometric distribution for modeling the enclosure experiment

For each polymorphic site *S*_i_ in the genotype data of the population we denote:

*N*: the number of valid genotypes at *S*_i_

**m**: the vector of absolute genotype frequencies before predation at *S*_i_, such that,

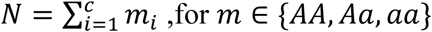

,where *c* is the number of possible genotypes

*n*: the number of mice successively removed from the enclosure by predators

**x**_p_: the vector of absolute genotype frequencies in the predated individuals at *S*_i_, such that,

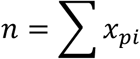

**x**_s_: the vector of absolute genotype frequencies in the predated individuals at *S*_i_, such that,

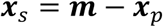

**w**: the vector of weights for the three possible genotypes at *S*_i_, where higher weights increase the chance of being predated.

The probability of observing a multivariate set **x**_p_, given **m** and **w** is given by the Wallenius’ non-central hypergeometric distribution (Chesson 1976, Fog 2008)

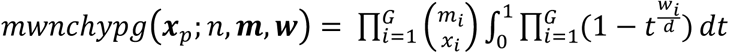

with 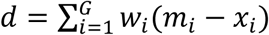

Given **x**_p_ and **m** it is also possible to estimate **w** using the function “oddsWNCHypergeo” in the R package “BiasedUrn” (Fog 2015)

The distribution of genotype frequencies in the set of survivors at the end of the experiment is given by the complementary Wallenius’ non-central distribution

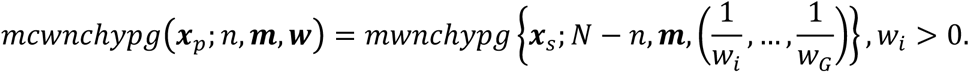

We will refer to 1/w_i_ as the fitness of genotype *i.*

## Constructing the null hypothesis that survival is independent of genotype at any locus

We can calculate for each variant, the probability that the distribution of genotype frequencies in the survivors is a random sample from the initial population. To do this we use the multivariate Wallenius non-central distribution with all weights set to 1. (In this special case, this distribution converges to the multivariate hypergeometric distribution).

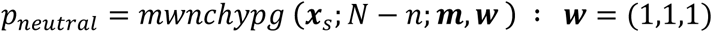

For each polymorphic site, the elements of **x**_s_ (*x*_s,AA_; *x*_s,Aa_; *x*_s,aa_) are calculated as:

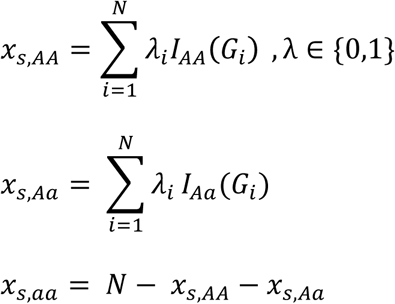

where λ is the survival vector such that λ *∈* {0,1} where 1 indicates survival, **G** is the vector of genotypes where **G** *∈* {AA, Aa, aa}, and *I*_AA_ is the indicator function such that *I*_AA_(*G*)=1 if *G*=“AA”.

This is the probability represented in Figure 4 (using a log10 transformation). In this figure, variants are categorized into significant and non-significant variants based on an arbitrary significance threshold of α=0.01. We define *Y*_obs_ as the number of significant variants in a subset of the total number of variant *S* such that,

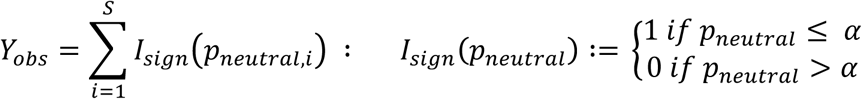

we then approximate the expected distribution of *Y* under the assumption that survival is independent of genotype by randomly permuting the survival vector λ 1000 times and recalculating *Y* for each of the permutated dataset. We let **Y**_neutral_ be the vector containing the 1000 random values of *Y* and calculate the p-value of *Y*_obs_ as:

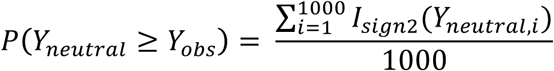

where,

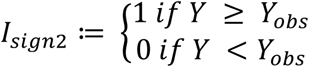

## Constructing the null hypothesis that survival depends only on a single locus under selection

In the cases where *Y*_obs_ is significant, we further consider the possibility that survival can be affected by a single site under selection *S*_sel_, where *S*_sel_ belongs to {1,2,…,*S*_max_}, the set of all variants in the dataset.

For each variant we calculate the fitness of each genotype using the multivariate Wallenius non-central distribution as explained above. We then obtain the selection and dominance coefficient *s* and *h* from the fitness values as

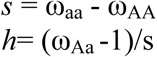

We set ω_aa_ to be the highest fitness among homozygote genotypes such that *s* is always positive or equal to zero and, when possible, we report whether the *a* allele corresponds to the ancestral or derived state (*an outgroup sequence was available for Agouti, but not for the background loci*).

We then calculate the likelihood ratio statistic *D* as,

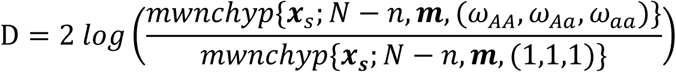

p-values for the test statistics are calculated under the assumption that *D* follows a chi-square distribution with 2 degrees of freedom.

We define *S*_sel_, the site under selection, as the site associated with the smallest P value.

To test whether *Y*_obs_ is significant under the assumption that survival depends on genotype at locus *S*_sel_ we use the same resampling approach described above, but we sample the survivors according to the multivariate non-central Wallenius’ distribution with the specific weights estimated for *S*_sel_. This can be done with the function rMWNCHC() in the R package “BiasedUrn” (Fog 2015). (Null distributions can be seen in Figure 5).

